# A major histocompatibility complex (MHC) class II molecule that binds the same viral pathogen peptide with both nonamer and decamer core sequences for presentation to T cells

**DOI:** 10.1101/2025.10.27.684722

**Authors:** Anastasia Goryanin, Atlanta G. Cook, Shahriar Behboudi, Jim Kaufman, Samer Halabi

## Abstract

Classical molecules encoded by the major histocompatibility complex (MHC) are central to immune responses. Compared to typical mammals, the chicken MHC is small and simple, determining life or death from economically-important pathogens like avian influenza virus and Marek’s disease virus (MDV). Several genes within the tightly-linked chicken MHC have been suggested to determine resistance and susceptibility to MDV, but it was a surprise to find that the dominantly-expressed class II molecule from the resistant B2 haplotype employed a novel peptide-binding mode with a decamer core sequence compared to the susceptible B19 haplotype with a typical nonamer core. We examined the crystal structure of the dominantly-expressed class II molecule from another resistant haplotype, B21, which is extremely frequent in commercial chicken flocks, to find that it bound the same MDV peptide with both nonamer and decamer cores, revealing an unexpected plasticity of binding that potentially increases the immune response to this devastating pathogen.

## Introduction

The major histocompatibility complex (MHC) of typical mammals like humans and mice is a large genetic region containing hundreds of genes with diverse functions, among which are a few highly polymorphic classical class I and class II genes that encode molecules which were discovered because of graft rejection. These class I and class II molecules bind peptides within cells and present them on the cell surface to thymus-derived (T) cells and natural killer (NK) cells, and therefore play crucial roles in the adaptive and some innate responses (1). With multigene families of class I and class II molecules, the human MHC determines relatively weak genetic associations with infectious diseases, especially compared to autoimmune diseases (2, 3).

In contrast to such typical mammals, the MHC of chickens is renowned for strong genetic associations with economically-important pathogens such as the herpesvirus that causes Marek’s disease (MDV), the orthomyxovirus that causes avian influenza, the acute retrovirus that causes Rous sarcomas and the coronavirus that causes infectious bronchitis (2–4). MDV has a complex life cycle and, while mostly controlled by vaccination (with billions of doses every year), is still a major problem for the global poultry industry (5, 6). The chicken MHC is only part of a larger B locus, which was discovered as a blood group and then associated with resistance to Marek’s disease during the intensification of the poultry industry (4, 7). However, resistance is associated with the chicken MHC (previously called the BF region or the BF-BL region) (8), which is notably small and simple, with dominantly-expressed classical class I (BF2) and class II B (BLB2) genes (2, 9–11).

Several genes have been suggested to account for resistance to Marek’s disease, including the dominantly-expressed class I gene (BF2), the dominantly-expressed class II B gene (BLB2), an NK receptor gene (BNK) and a gene of unknown function called BG1 (12–15). However, it has not been possible to establish which candidate gene is responsible, since no recombination within the chicken MHC has been directly observed in deliberate matings (16–19), although historic recombinants have been reported (18, 20).

One clue to a possible resistance mechanism is the discovery that the dominantly-expressed class II molecule from the MDV-resistant MHC haplotype B2 binds antigenic peptides in an unprecedented way compared to all known class II molecules (14, 21), including to one from the MDV-susceptible haplotype B19 (22). All previously-described class II molecules bind nonamer core peptides in an extended conformation along the peptide-binding groove, with sequences extending out of the groove on both ends (23). However, BL2*02:01 (composed of the monomorphic α chain BLA and the β chain from the B2 haplotype, BLB2*02:01) binds peptides (including multiple antigenic peptides from MDV) with a decamer core.

Comparison of the crystal structure of BLB2*02:01 with other structures revealed a crinkle in the peptide backbone caused by one smaller MHC residue within the groove which meant that ten amino acids were required to span the groove rather than nine (14).

We wished to determine whether this difference in core peptide length between BL2*02:01 from a disease-resistant haplotype and BLB2*19:01 from a disease-susceptible haplotype might also be true for other MHC haplotypes. Using enzyme-linked immunosorbent spot (ELIspot) assays to assess T cell responses from birds with the MDV-resistant haplotype B21, a peptide was identified to which the CD4 cells of all B21 birds responded strongly (24, 25). Upon peptide assembly and X-ray crystallographic analysis, we found to our surprise that the BL2*21:01 binds this peptide in both the nonamer and decamer conformations, potentially stimulating more T cells and leading to a greater protective immune response.

## Materials and Methods

### Cell lines

Sf9 insect suspension cells (*Spodoptera frugiperda* female ovarian cell line, ATCC CRL-1711) were used for production of baculovirus, and High Five insect suspension cells (*Trichoplusia ni* female ovarian cell line, GIBCO B85502) were used for protein production, both grown in Insect-XPRESS Protein-free Insect Cell Medium (Lonza, BE12-730Q) supplemented with L-glutamine (to 1%), 50 U/ml penicillin, and 50 μg/ml streptomycin at 27°C with shaking at 135 rpm.

### Cloning, protein expression, and purification

For protein production and crystallography, the different constructs were cloned separately into the baculovirus insect cell expression vector, pFastBac1 (GIBCO 10360014), modified to contain an N-terminal GP67 secretion signal sequence (Supplementary Fig. S1). For the BLA construct, the GP67 signal sequence was followed by the ectodomain of the α-chain of BLA (Hα5 to Eα188; GenBank accession number AY357253 (but with S79 due to dimorphism noted in (26), then a short Gly/Ser-linker (G4S), a c-fos dimerization domain (27), a two amino acid linker (GT), and an Avi-tag sequence for biotinylation. For the BLB2*2101 construct, the GP67 signal sequence was followed by a peptide sequence (Supplementary Fig. S1), a 15 amino acid Gly/Ser-linker [3(G_4_S)], the ectodomain of the β-chain of BLB2*2101 (Fβ6 to Kβ198; DQ008585.2/AB426152.1), a TEV-protease cleavage site, a c-jun dimerization domain (27), a Gly/Ser-linker [2(G_4_S)], a V5-tag, and a 10x His-tag. Correct cloning was confirmed by DNA sequencing.

To generate bacmids, the constructs were transformed into DH10Bac cells (GIBCO 10361012), and the integrated genes into bacmids were extracted from white colonies following X-gal selection. The extracted bacmid DNA was subjected to PCR using M13 primers to confirm transposition of inserts, and then transfected into Sf9 cells to generate P0 viral stocks. First, 20 μl bacmid DNA and 15 μl Fugene (Promega E2311) were each diluted into 600 μl of Lonza Insect Xpress medium, mixed, and incubated at room temperature for 20 minutes, and then 200 μl of the mixture were transferred to each well of a 6-well plate containing 9 × 10^5^ Sf9 cells in 2 ml antibiotics-free medium and incubated for 4 to 6 days at 27°C. The cell medium containing baculovirus was collected, passed through a 0.2-μm filter (Sartorius 16532-K), and then this P0 stock was used to amplify the number of viruses to give the P1 stock as follows: 1 ml P0 stock was added to 50 ml of 2 × 10^6^ Sf9 cells/ml in suspension and cultured at 27°C for 48 hours with shaking at 135 rpm, then the cells were pelleted, and the culture media was filtered as above. Both P0 and P1 viral stocks were stored at 4°C until required.

High Five cells were grown in 0.6 L medium in 2 L flasks and transduced with 6 to 10 ml P1 stock. Forty-eight to seventy-two hours after infection, culture media was collected and incubated overnight at 4°C with nickel-Sepharose Excel (GE Healthcare 17-3712-01). The His-tagged protein complex was eluted from the nickel-Sepharose with 1 M imidazole in 25 mM Tris-HCl (pH 8.5), then purified by size-exclusion chromatography on a fast protein liquid chromatography (FPLC) instrument (Amersham) using Superdex S200 in 100 mM TrisCl (pH 8.5). Then the purified protein complex peak was subjected to endoproteinase Glu-C (V8 Protease, Sigma, United Kingdom) cleavage at 37°C overnight to remove the C-terminal tags and the dimerization domains, with successful proteolysis confirmed by size shift in SDS-PAGE. The proteins were again purified as above using Superdex S200 in 25 mM TrisCl (pH 8.5), and then concentrated using Amicon Ultra (with 3,000 molecular weight cut-off, Merck UFC8003) to 3 to 12 mg/ml as determined by a nanodrop spectrophotometer.

### Crystallization and structure determination

Crystallization conditions were screened using the PEGs II suite (Qiagen, United Kingdom) at 20°C, with 11 mg/mL protein in 25 mM TrisCl (pH 8.5) mixed at 1:1 or 1:2 with mother liquor to give 0.6 to 0.9 μl sitting drops. The protein complex BL2*021:01 with the peptide AVVHSVRALMLAERQ from MDV phosphoprotein 38 (pp38 residues171-186) was crystalized in 100 mM sodium acetate pH 4.6 with 15 % w/v PEG 4000. Crystals were cryo-cooled in mother liquor supplemented with 35% (v/v) glycerol before data collection.

Diffraction data were collected remotely on the I04-1 beamline (Diamond Light Source, Oxford, UK) at a wavelength of 0.920 Å. Data reduction and scaling were performed using xia2 (28) and AIMLESS (29). The crystal of the BL2*021:01 belongs to the P3_2_21space group, and the structure was solved by basic molecular replacement deploying Phaser from the CCP4i package (30) using BL2*019 (6KVM) as the search model (22). Further rounds of manual model building and refinement were done using Coot (31) and REFMAC5 (32). Further details about collection and refinement (Supplementary Table S1) were generated using Phenix (33). The structure was deposited in the Protein Database (PDB) on 9 August 2021 and assigned accession number 7PDY.

### Interaction analysis

The online software, Protein Data Bank in Europe Proteins, Interfaces, Structures and Assemblies (PDBePISA) (34) was used to calculate the potential interactions from the crystal structure, and their crystal contacts. PyMOL (The PyMOL Molecular Graphics System, Version 2.0 Schrödinger, LLC) was used to display and analyze the structural data visually.

### Differential scanning fluorimetry

Differential scanning fluorimetry was performed using a CFX Connect real-time PCR thermal cycler (BioRad, United Kingdom) set to Fluorescence Resonance Energy Transfer (FRET) scan mode. SYPRO orange (Invitrogen S6651) was diluted to 1:500 in 25 mM TrisCl (pH 8) and 12.5 μL mixed with 12.5 μL 10 μM protein in 25 mM TrisCl (pH 8) in each well of a 96-well plate, with buffer alone serving as negative control. The samples were scanned every 0.5°C from 25 to 95°C. Data analysis was performed in Prism (GraphPad, San Diego, California, USA).

## Results

### The dominantly-expressed class II molecule from the B21 MHC haplotype binds the MDV pp38 peptide

A single peptide (AVVHSVRALMLAEARQ) from the MDV-encoded phosphoprotein 38 (pp38, residues 171-186 from the very virulent MDV strain RB1B) was found to strongly stimulate CD4 cells of all vaccinated and/or infected B21 birds, as assessed by ELIspot, cytokine and proliferation assays (24, 25). As was previously constructed for several MDV peptides bound to BL2*002:01 (14), a nucleic acid sequence encoding this pp38 peptide was inserted between a signal sequence and the β chain of the dominantly-expressed class II molecule of the B21 haplotype, BL2*021:01, which was then expressed in insect cells together with the monomorphic chicken class II α chain (BLA). The α and β chain sequences were both followed by a linker with an endoproteinase V8 cleavage site preceding c-fos or c-jun sequences designed to promote association of the two chains (Fig. 1A, Supplementary Fig. 1).

**Figure 1.**
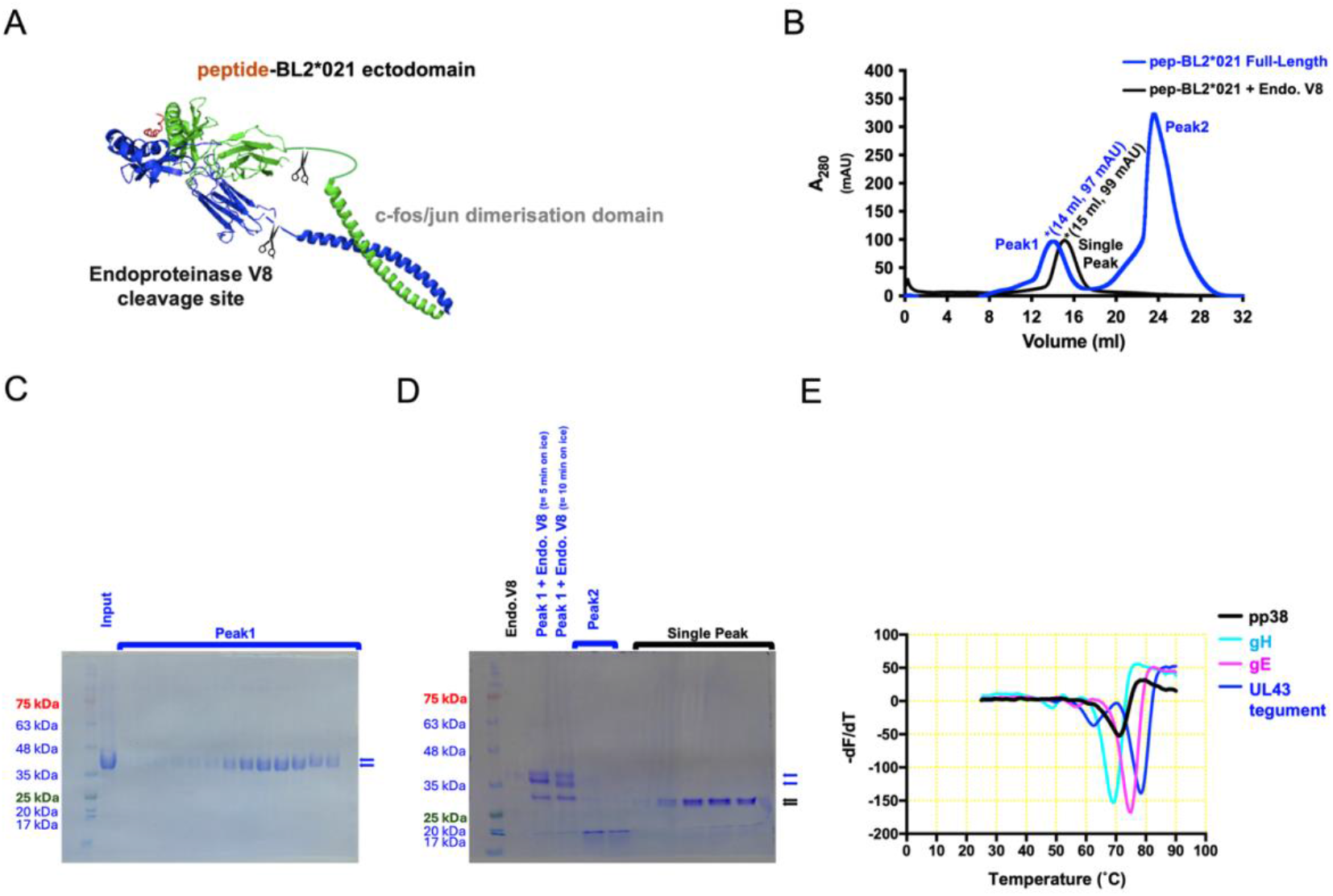
BL2*21:01 linked to the MDV pp38 peptide was expressed in insect cells and purified, and has a similar thermostability as other chicken class II molecules bound to MDV peptides. A. Cartoon showing full-length construct, with scissors indicating the potential cleavage sites of endoproteinase Glu-C (Endoproteinase V8). B. UV traces of FPLC size-exclusion chromatography (SEC) using Superdex S200 column, before (blue) and after (black) being subjected to Endoproteinase V8 cleavage at 37°C overnight to remove C-terminal tags and dimerization domains. C. SDS-PAGE followed by Coomassie blue staining after initial nickel column purification (Input) and SEC fractions comprising Peak1 (blue trace of profile in A). Standard protein markers with indicated molecular masses; blue arrows indicate position of α- and β-chains. D. SDS-PAGE followed by Coomassie blue staining of endoproteinase V8 on its own, Peak1 (blue trace of profile in A), and SEC fractions from Single Peak (black trace). Standard makers and blue arrows is in C, black arrows indicate the position of the endoproteinase V8 cleaved α- and β-chains. E. Thermal denaturation curves for BL2*02:01 molecules expressed with MDV peptides from glycoprotein H (gH, with melting temperature 69°C), glycoprotein E (gE, 75°C), and unique long gene 43 tegument protein (UL43, 78°C), compared to BL2*021:01 molecule expressed with MDV pp38 peptide.

Successful expression shows that this immunogenic pp38 peptide does indeed bind the dominantly-expressed class II molecule of the B21 haplotype (Fig. 1B, C). Proteolytic removal of the fos/jun dimerization sequences led to a smaller but highly pure monomer corresponding to the extracellular domains of the class II heterodimer bound to peptide (Fig. 1B, D). A thermostability assay showed that the pp38 peptide bound to BL2*021:01 has a melting temperature around 70°C, in the same range as the MDV peptides bound to BL2*002:01 (Fig. 1E).

### The overall structure of the dominantly-expressed class II molecule from the B21 MHC is similar to other class II molecules

The purified monomer of BL2*021:01 bound to the pp38 peptide was crystallised successfully, diffraction data were collected from the Diamond Light Source synchrotron, and the structure was solved by molecular replacement using BL2*019 (6KVM) as the input model, followed by rounds of manual model building and refinement. The crystals were found to belong to the space group P3_2_21, with unit cell dimensions a, b and c being 238.5, 238.5 and 76.4 Å, respectively; with angles α, β and γ being 90°, 90° and 120°, respectively (Supplementary Table S1).

The asymmetric unit was found to contain three monomers. The monomers composed of C/D chains and E/F chains are oriented with their peptide-binding grooves (PBGs) facing away from each other, with the PBG of the C/D monomer facing the PBG of the A/B monomer (Fig. 2A). Superposition of the three monomers showed that their structures are overall very similar (Fig. 2B), with root mean square deviations (RMSD) of the alpha carbons (Cα) being less than 0.4 Å (Fig. 2E).

**Figure 2.**
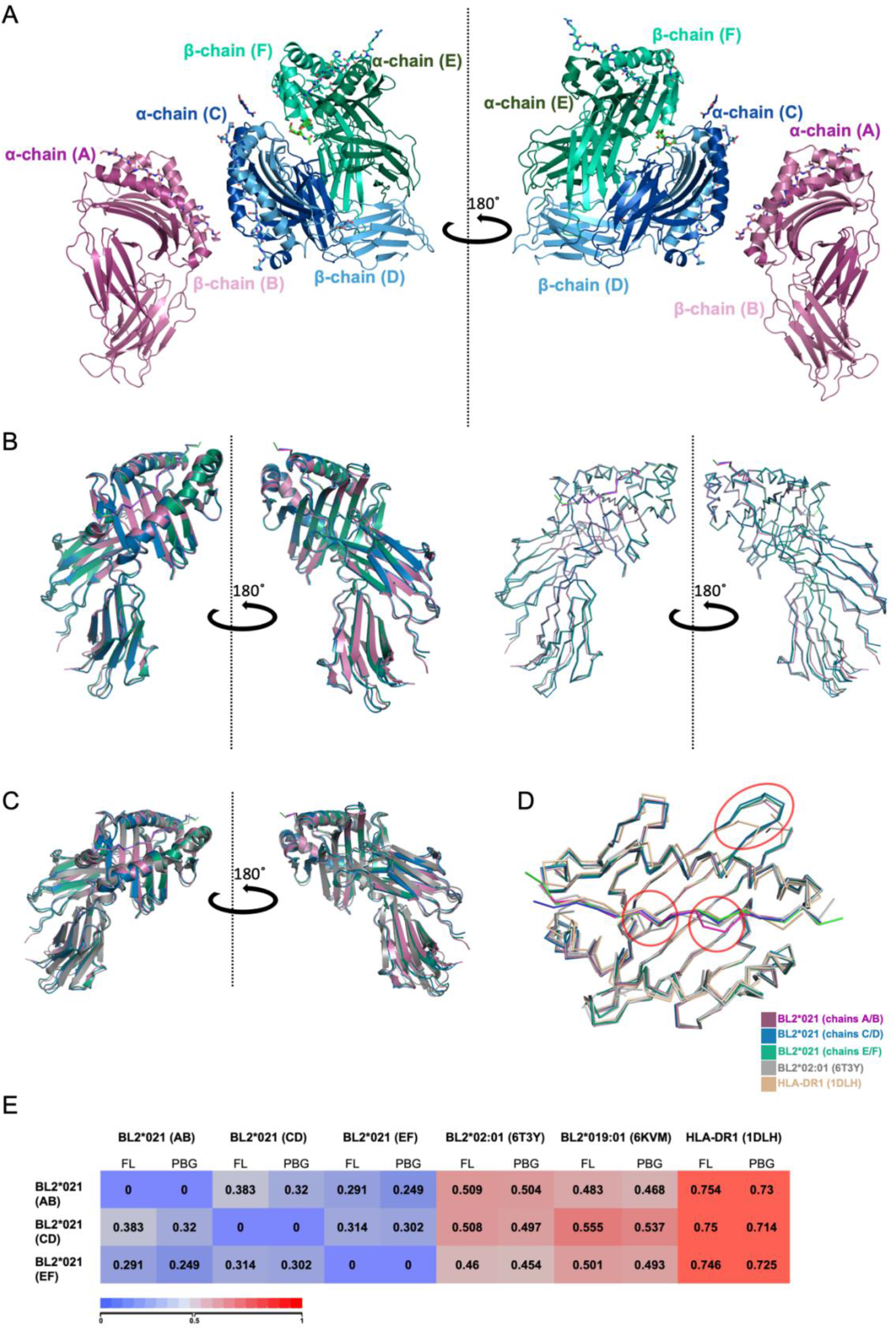
Three molecules of BL2*21:01 linked to the pp38 peptide were found in the asymmetric unit (7PDY), with overall structures similar to other class II molecules. A. Two side views rotated by 180° around the y-axis of BL2*021:01 in PyMol cartoon mode and peptide in Pymol ribbons style; chains A and B in dark and light magenta (bound peptide in light pink); chains C and D in dark and light blue (bound peptide in tv-blue), chains E and F in dark and light green (bound peptide in tv-green). B. the three BL2*021:01 monomers in the asymmetric unit overlayed in top of each other in cartoon (left) and ribbons (right). C. As in B but including HLA-DR1*01 (PDB ID 1DLH, dark grey). D. Top view of the peptide-binding groove (PBG) in ribbons (that is, Cα backbone) for the three monomers of BL2*21:01 linked to the pp38 peptide in the asymmetric unit, along with BL2*02:01 (6T3Y, grey) and HLA-DR1*01 (1DLH, orange). Significant differences are circled. E. RMSD values comparing the Cα backbone of the full length (FL) molecules as in A and their peptide binding grooves (PBG); scale from 0 (blue) indicating identity to 1 (red) indicating lower similarity.

Comparison of the three monomers from B21 with structures from the chicken MHC haplotypes B2 and B19 (14, 22), as well as the human molecule HLA-DR1*01:01 (35), again showed great similarity (Fig. 2C, D), the RMSD with the chicken alleles being less than 0.56 Å and with the human molecule being less than 0.76 Å (Fig. 2E). There are three notable differences (features circled in Fig. 2D). The length of one loop of the chicken α chain (residues 16-19 of the mature protein) is four amino acids longer than in humans and other mammals (22, 26). The peptide bound to BL2*002:01 has a crinkle leading to a decamer core sequence (14), and the peptide bound to one of the BL2*021:01 monomers also has a crinkle in the peptide.

### Two B21 monomers bind the pp38 peptide with a canonical nonamer core sequence, but one binds with a decamer core sequence

Almost all class II molecules are reported to bind peptides with nine residues (the nonamer core sequence) that lie flat as a poly-proline II helix in the PBG, with several class II residues contacting peptide mainchain atoms for stability, typically with four peptide sidechains inserting into pockets for specificity, and with the peptide extending out as flanking residues on either end of the PBG (23, 35–37) (Fig. 3). The exception has been the dominantly-expressed class II molecule of the B2 haplotype, BL2*002:01, which was unexpectedly found to bind peptides with ten amino acids in the PBG to give a decamer core sequence (14) (Fig. 3C). Given that the MDV-susceptible B19 haplotype has a class II molecule with a canonical nonamer core (22) (Fig. 3B), it seemed possible that the MDV-resistant B2 and B21 haplotype might both have class II molecules with the unusual decamer core.

**Figure 3.**
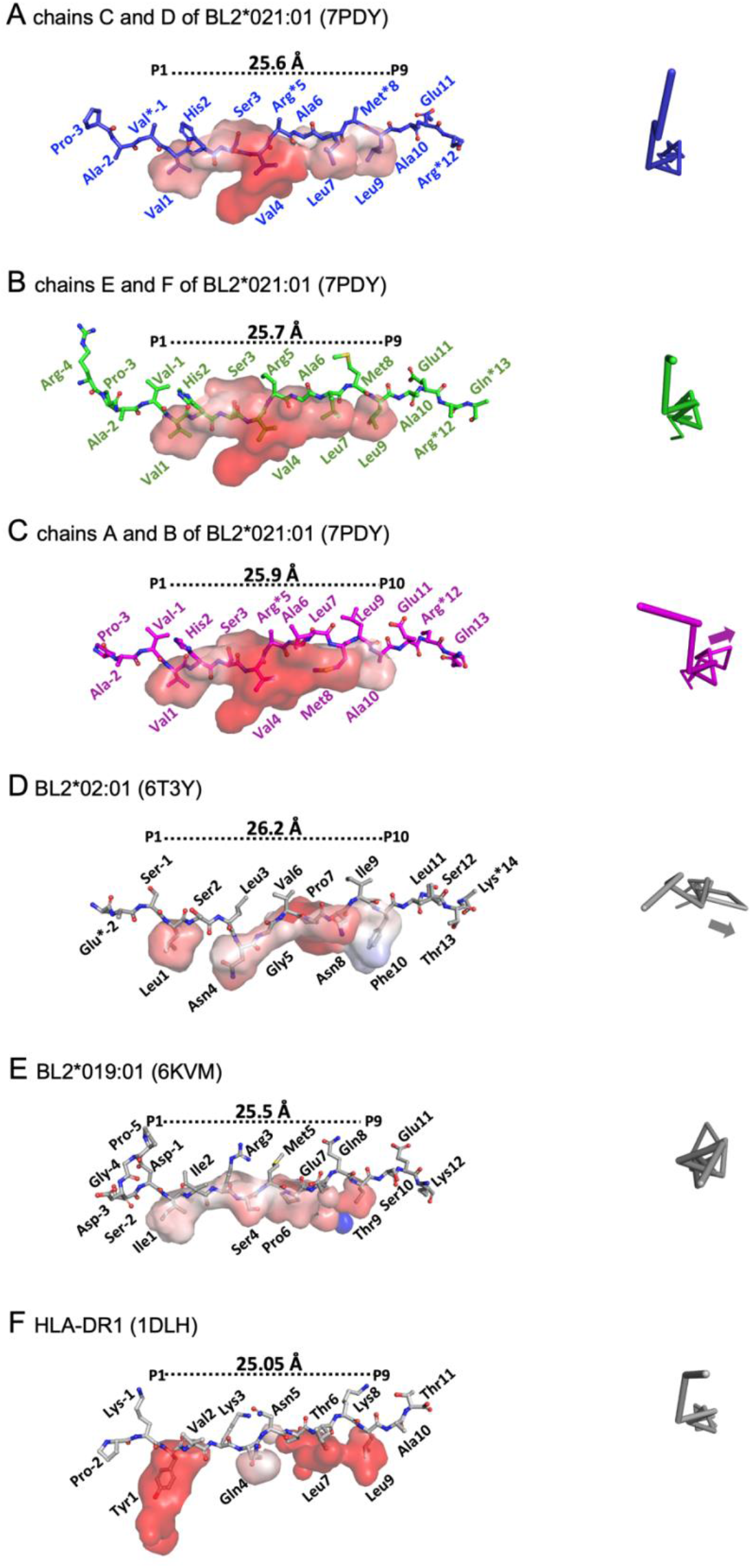
Two molecules of BL2*21:01 in the asymmetric unit bound the pp38 peptide with a typical nonamer core sequence, but one bound a decamer core sequence with peptide positions P6 and P7 departing from the typical polyproline II helix. Panels A-F. Left, side view of peptide in PyMol sticks with amino acids indicated, length between Cα of the first and last amino acid in the core sequence indicated, and pockets as surfaces. Right, Cα of the bound peptide in PyMol ribbons viewed down the axis of the polyproline helix, with arrows indicating departure from helix. A. BL2*21:01 C/D monomer (7PDY) nonamer, B. BL2*21:01 E/F monomer (7PDY) nonamer, C. BL2*21:01 A/B monomer (7PDY) decamer and arrow, D. BL2*02:01 (6T3Y) decamer and arrow, E. BL2*19:01 (6KVM) nonamer, F. HLA-DR1*01 (1DLH) with nonamer core sequence.

In fact, the B21 A/B monomer in the asymmetric unit does have a decamer core (AVVHSVRALMLAEARQ with core underlined) like B2 (lefthand panels of Fig. 3C, D), but the C/D and E/F monomers have nonamer cores (AVVHSVRALMLAEARQ with core underlined) like B19 and the human class II molecule HLA-DR1 (lefthand panels of Fig. 3A, B, E, F). The peptides are in polyproline II helices, including along most of the peptide bound to the B21 A/B monomer and to the B2 monomer, except that each of these two peptides has a crinkle in which a few residues depart from the helix (arrows in the righthand panels of Fig. 3C, D).

As a result of these crinkles, the C-terminal portions of the peptides are pulled in the N-terminal direction, leading to a shift in peptide residues that fit into the pockets. The length of the binding site remains roughly the same for all of the PBGs, but the pockets shift from binding the sidechains of peptide positions P_1_, P_4_, P_6_ and P_9_ for B19, and P_1_, P_4_, P_7_ and P_9_ for DR1 and the B21 C/D and E/F monomers, to P_1_, P_4_, P_8_ and P_10_ for both B2 and the B21 A/B monomer (lefthand panels of Fig. 3).

The crinkles also change the register of the mainchain atoms of the peptide that make hydrogen bonds (H-bonds) with the class II residues (Fig. 4A). Most of the H-bonds originally described for HLA-DR1 (35) are conserved for the B21 C/D and E/F monomers, and for the N-terminal part of the B21 A/B monomer. However, after peptide position 6 (P_6_), the H-bonds for the B21 A/B monomer are offset by one amino acid. Thus, Trpβ61 contacts the P_8_ carbonyl of the peptide in DR1 and the two B21 monomers but it contacts the P_9_ carbonyl of the B21 A/B monomer, Asnα69(73) contacts the P_9_ amino group compared to the P_10_ amino group, Argα76 (Nα80) contacts the P_10_ carbonyl group and the P_12_ amino group compared to the P_11_ carbonyl group and the P_13_ amino group, and Aspβ57(Qβ57) contacts the P_10_ amino group and carbonyl group compared to the P_11_ amino group and carbonyl group.

**Figure 4.**
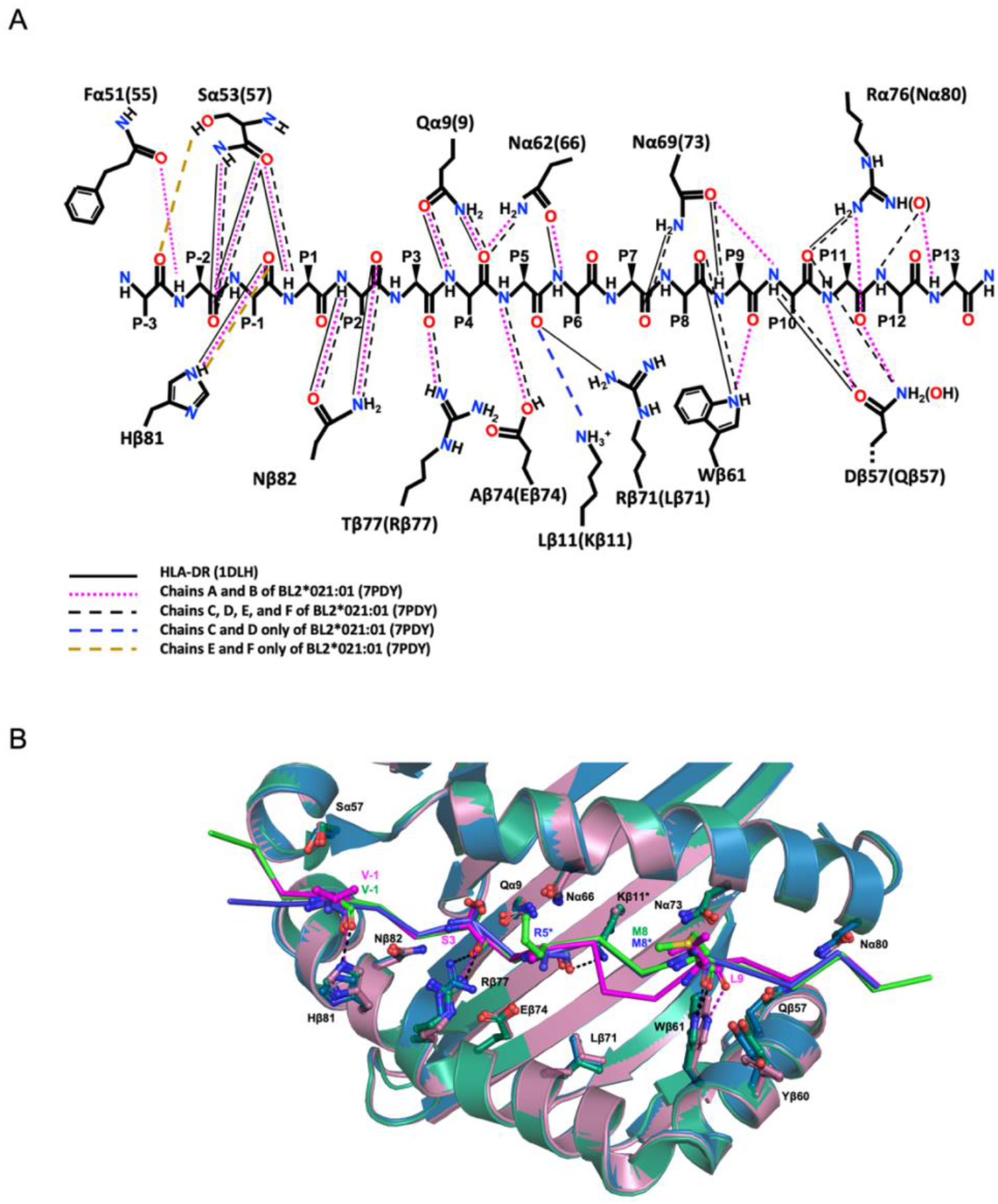
The H-bond network between BL2*21:01 and the pp38 peptide with a decamer core sequence shifts register at P6, but this is not due to changes in the conformation of class II residues. A. Shift at P6 shown in schematic of H-bonds interacting with peptide main chain atoms (cut-off of 4 Å) originally described for HLA-DR1*01 (solid black lines) compared to BL2*021:01; chains A and B (dotted pink lines), chains C, D, E, and F (dashed black lines), chains C and D (blue dashed lines), and chains E and F (orange dashed line). B. Different rotamers and slight differences in orientation of class II amino acids found between the three monomers in the asymmetric unit did not significantly affect H-bonds. Superimposition of the PBG of BL2*021:01 in PyMol cartoon mode with bound peptides in Pymol ribbons style; chains A and B in magenta, with bound peptide in light pink; chains C and D in blue, with bound peptide in tv-blue, chains E and F in green, with bound peptide in tv-green. The side chains of amino acids that show some deviation are shown in PyMol sticks mode and are named in black. Hydrogen bonds that formed between these side chains and the peptide are indicated by dotted lines as in panel A.

In these conformational rearrangements (Fig. 4A), H-bonds are gained between the Lysβ11 amino group and the P_5_ carbonyl group in C/D and E/F (but not A/B, and replacing Argβ72 in HLA-DR1), and lost between the Asnα62(66) carbonyl group and the P_6_ amino group in the C/D and E/F monomers, as well as between the Asnα69(73) amino group and the P_7_ carbonyl group in the A/B monomer. Due to species differences in amino acid sequence, H-bonds are gained in all B21 monomers compared to HLA-DR1 between the Argβ77 guanidino group and P_3_ carbonyl group, the Gluβ74 carboxy group and the P_5_ amino group, the Asnα62(66) amino group and the P_4_ carbonyl group, the Glnβ57 amino group with either the P_10_ carbonyl of C/D and E/F or the P_11_ carbonyl of A/B, and the Asnα80 amide carbonyl group with either P_12_ amino group of C/D and E/F or the P_13_ amino group of A/B.

### The two conformations of peptide bound to BL2*021:01 reveal a plasticity of binding

Since the chicken class II α chain (BLA) is not polymorphic (26), any differences seen in peptide-binding motif must be due to the β chain (BLB) (9, 20), which is highly polymorphic in the β1 domain that forms half of the PBG but not in the invariant immunoglobulin-like (Ig-like) β2 domain or the connecting peptide (CP) (Fig. 5). The cause of the crinkle that leads to a decamer core for the dominantly-expressed class II molecule from the B2 haplotype is clear. Instead of the relatively large sidechain at position β72 found almost all class II molecules (Phe in B19, Tyr in HLA-DR1), BL2*002:01 has the relatively small sidechain of Ser, which draws the Asn at P4 downwards and thus pulls the C-terminal portion of peptide towards the N-terminus (Figs. 5, 6A).

**Figure 5.**
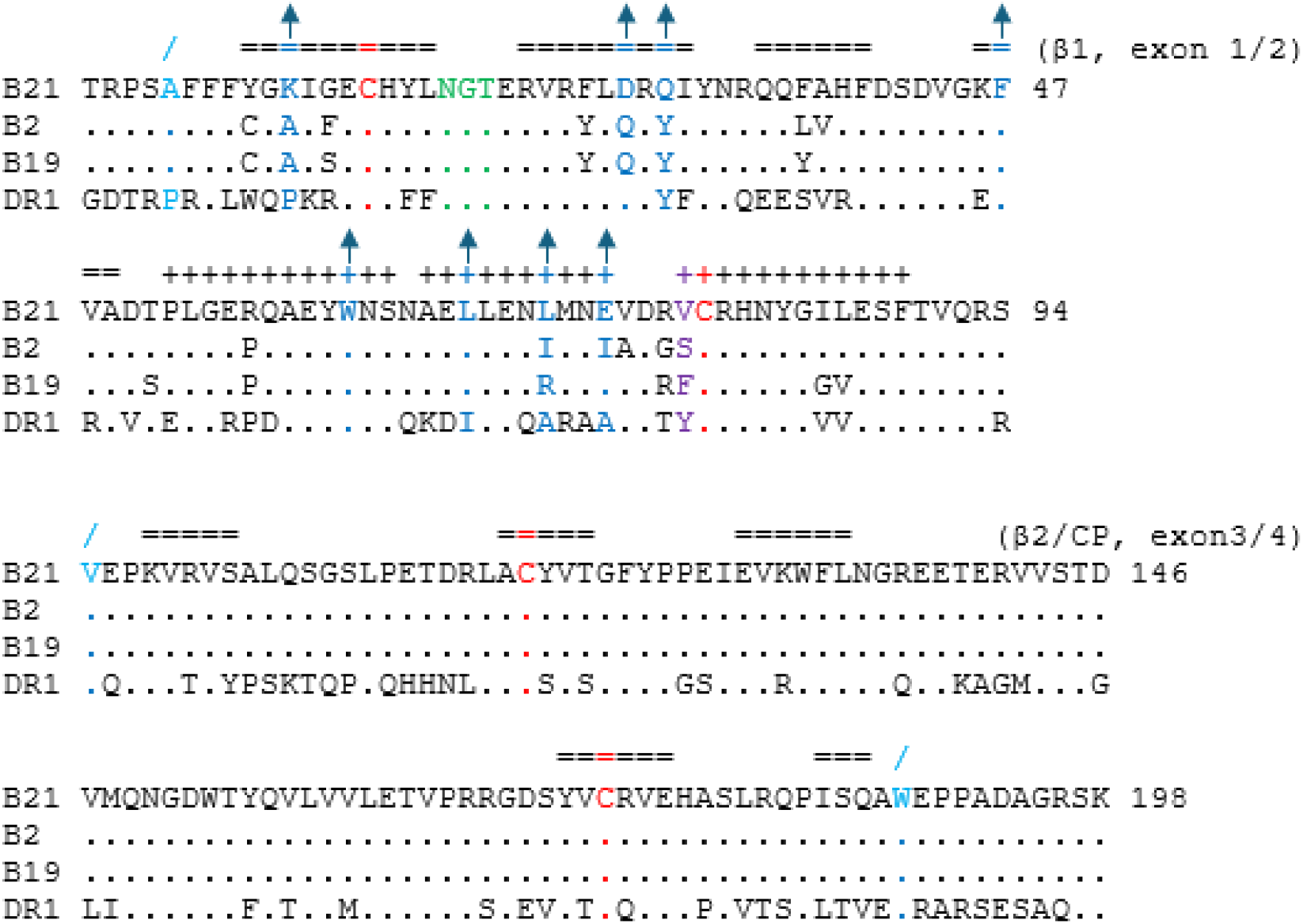
Alignment of β chain sequences shows that polymorphism in chicken class II molecules is found in the β1 domain, and that the amino acids contacting the bulge of the pp38 peptide in the decamer core are mostly different in BL2*21:01 and other chicken class II molecules. Amino acid sequences of the extracellular domains of β chains of BL2*21:01, BL2*02:01, BL2*19:01 and HLA-DR1*01 were aligned, starting with the first amino acid of the mature chain. Amino acids at intron/exon boundaries are indicated in light blue with a slash above, cysteines involved in the intradomain disulphide bonds in red, glycosylation site for N-linked glycan in green, the amino acid at position 78 responsible for peptide crinkle in BL2*02:01 and secondary structure symbols in purple, and amino acids and secondary structure symbols near the departure from the polyproline helix in the BL2*21:01 A/B molecule in blue, with arrows above also in blue. β1, β1 domain; β2, β2 domain; CP, connecting peptide. Single letter code, sequences and secondary structure (=, β sheet; +, α helix) from PDB files 7PDY, 6T3Y, 6KVM and 1DLH.

**Figure 6.**
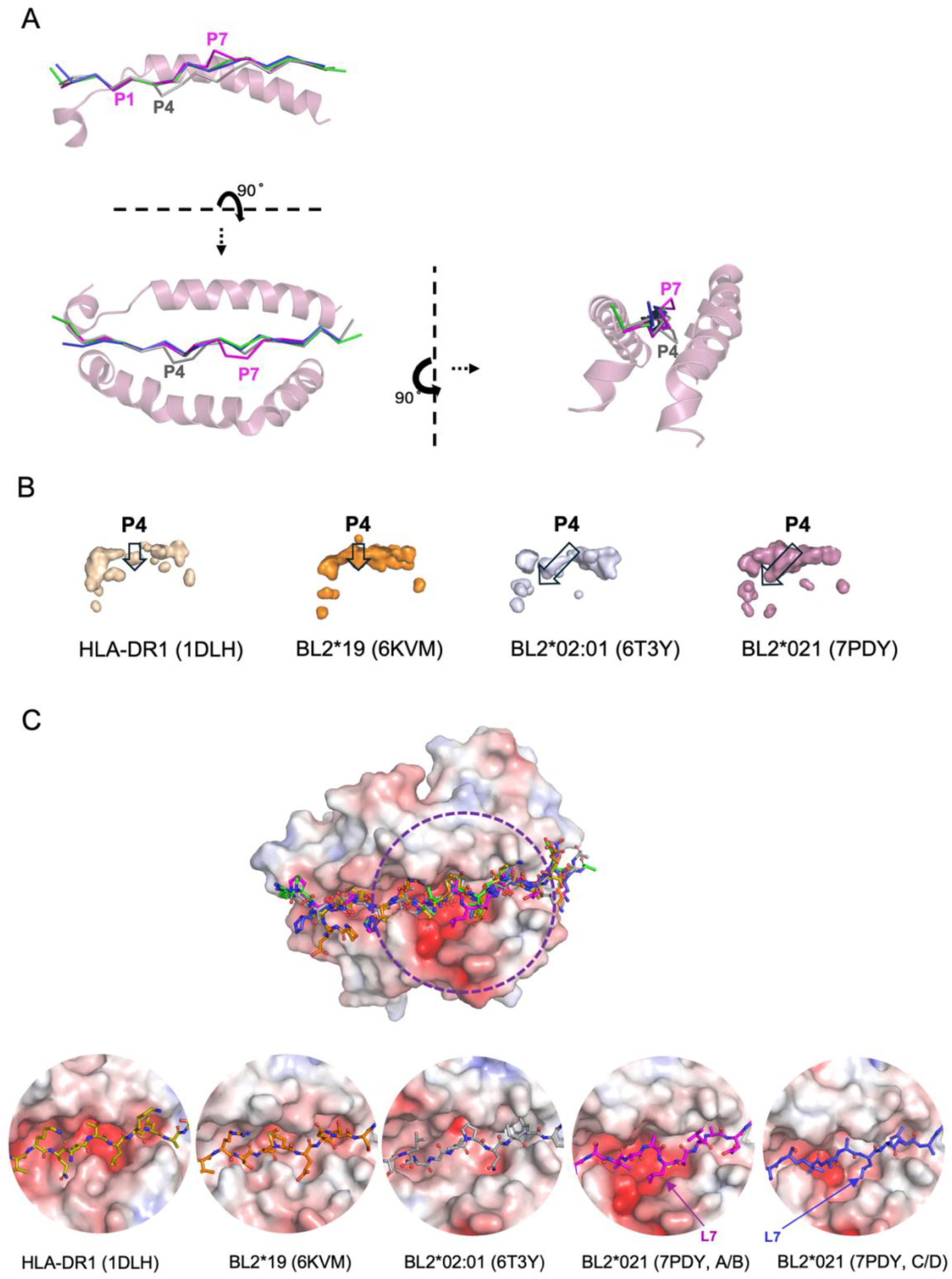
The departure from the typical polyproline II helix in the decamer core sequence is located at position P7 of the peptide bound to BL2*21:01 compared to P4 in BL2*02:01, but correlates with the size of pocket at P4 in both decamer peptides. A. Side, top and end-on views of the PBG in PyMol cartoon mode and the peptide in PyMol ribbons (Cα backbone) style, showing that the peptides in the C/D (tv-blue) and E/F (green) monomers of BL2*21:01 are in the typical polyproline helix, but depart from the helical conformation to the side and upwards at P7 for the A/B monomer of BL2*21:01 (7PDY, magenta) and to the side and downwards at P4 for BL2*02:01 (6T3Y, grey). B. Side view of PBG with pockets shown as surfaces, with APBS electrostatics calculated by PyMol (red, negative; blue, positive), and with pocket at P4 being small and vertical in HLA-DR1*01 and BL2*19:01 structures, but large and slanted forward in BL2*02:01 and BL2*021:01 structures. C. The surface of the class II molecule underlying peptide position P7 in the A/B monomer of BL2*21:01 is similar to the C/D monomer and to BL2*02:01, but narrower in BL2*19:01 and HLA-DR1*01. Top panel, superimposition of peptides over the PBG of BL2*21:01; bottom panel, peptides in the PBG of HLA-DR1*01 (1DLH), BL2*19:01 (6KVM), BL2*02:01 (6T3Y), the A/B and C/D monomers of the BL2*21:01 (7PDY). APBS electrostatics of surfaces calculated by PyMol (red, negative; blue, positive).

The cause of the crinkle that leads to the decamer core for the B21 A/B monomer is less clear. In fact, BL2*021:01 has a small Val at β72 (Fig. 5), which leads to a larger pocket for P_4_ that slants downwards towards the N-terminus like that of BL2*002:01, rather than the smaller downward-pointing pocket of BL2*019:01 or HLA-DR1 (Fig. 6B). However, there is no crinkle at P_4_ in any of the B21 monomers. Instead, the peptide in the B21 A/B monomer deforms from P_7_, with the Cα followed by the peptide bond found up and out of the PBG (Fig. 6A). The amino acid side chains that contact the peptide in all three monomers of B21 are the similar (Fig. S2). Comparison of the sequences of the dominantly-expressed class II molecules of B2, B19 and B21 shows that the B21 has significant differences in the contact residues near the crinkle, including Lys11 in B21 (Ala11 in B2 and B19), Asp28 (Gln28), Gln30 (Tyr30), Leu71 (Ile71, Arg71) and Glu74 (Ile74, Glu74) (Fig. 5). Comparison of the structures shows that the surfaces of the PBG in this region also differ significantly between B21 and the other MHC haplotypes (Figs. S3, S4).

### The decamer core found in BL2*21:01 is not due to various potential artefacts

The fact that the decamer core is found in only one of the three monomers in the BL2*21:01 crystal raises the possibility that it is some kind of artefact of crystallization. We were able to eliminate three obvious possibilities.

First, it seemed possible that different rotamers of class II residues might lead to changes in the conformation of bound peptides. Superposition of the B21 monomers show that there is little difference between C/D and E/F, which differ only slightly from A/B at Tyrβ60, Trpβ61, Argβ77 and Hisβ81 (Figs. 4B, S2). The greatest deviation is for Tyrβ60 which does not contact the peptide; Trpβ61 simply shifts the H-bond by one amino acid, Argβ77 changes rotamer but loses only one of two H-bonds in the E/F monomer, and Hisβ81 may lose the H-bond to peptide position P_-1_ but only in the C/D monomer (Table S2). Therefore, there is no strong evidence for conformational changes in class II residues leading to conformational changes in peptide.

Second, it seemed possible that contacts between the monomers within the asymmetric unit might influence the conformation of the bound peptides, based on the apparently close apposition of the PBGs of the A/B and C/D monomers (Fig 2A). Although there are many contacts between the C/D and E/F monomers (Fig. S5), a detailed examination showed that there is only one contact between the A/B and C/D monomers, with no other close approaches of the two surfaces: His at P_-3_ before the peptide core sequence bound to the A/B monomer forms a H-bond with Glnα61 of the α chain of the C/D monomer, both located far away from the peptide in the core region (Figs. 7A, S5, Table S3). Thus, there is no evidence that interactions between monomers within the asymmetric unit are responsible for the decamer core.

**Figure 7.**
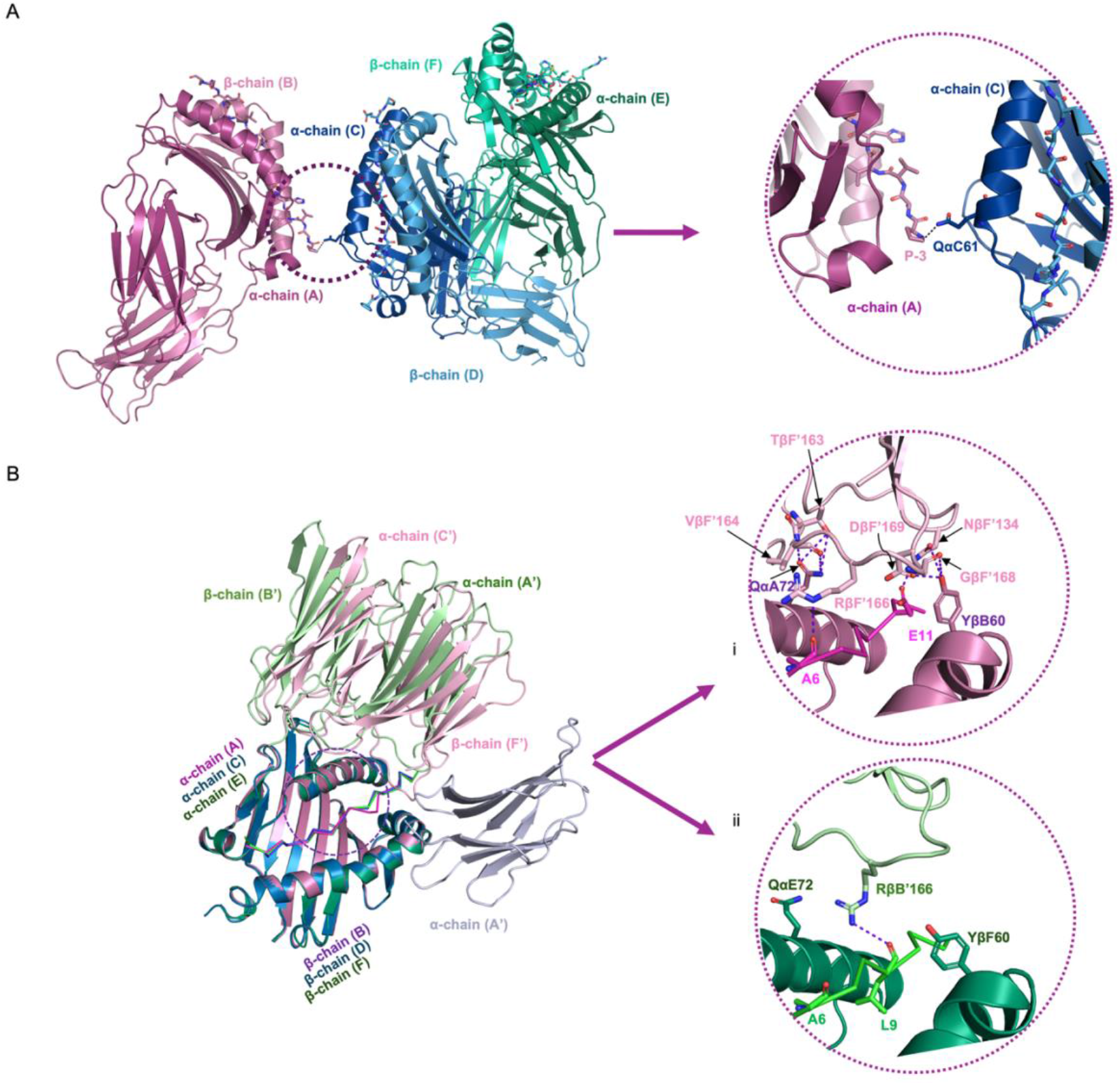
Contacts between monomers within the asymmetric unit and with monomers outside the asymmetric unit do not affect peptide binding in the decamer core. A. Interactions of A/B monomer with the two other monomers in the asymmetric unit are limited to a single contact involving part of the peptide far outside the PBG. His at P_-3_ from the A/B monomer forms an H-bond with Glnα61 from the C (α) chain. Colours as in Fig. 2A. B. The interactions between monomers within and outside of the asymmetric unit (that is, the crystal contacts) differ for A/B, C/D and E/F, but the near perfect superimposition indicates that there are no significant changes to the mainchain atoms. Argβ166 from the B’ and F’ chains both form H-bonds with mainchain carbonyls of the bound peptides, B’ after peptide Leu9 of E/F and F’ after peptide Ala6 of A/B, respectively. A/B monomer (PyMol cartoon mode, cb_magenta) and peptide (ribbon, light magenta) interacting with chains C’ and F’ (light pink); C/D (cb_blue) and peptide (tv_blue) with chain A’ (light blue); E/F (cb_green) and peptide (tv_green) with chains A’ and B’ (light green).

Third, it seemed possible that crystal contacts from molecules outside the asymmetric unit might influence the conformation of the bound peptides differently in the three B21 monomers within the asymmetric unit (Fig. S6). The molecules in adjacent asymmetric units do interact with the three B21 monomers within the asymmetric unit (Figs. 7B, 8), with the 4 Å proximity footprints of adjacent Ig-like domains showing minor contact on the β chain for C/D, more contact on the middle of the α chain and the C-terminal end of the peptide for E/F, and even more contact across the α chain, middle of the peptide and a bit on the β chain for A/B (Fig. 8). Superimposition shows that the interactions of the adjacent Ig-like domains are similar but rotated slightly for E/F and A/B, while being completely different for C/D (Fig. 7B, Table S4).

**Figure 8.**
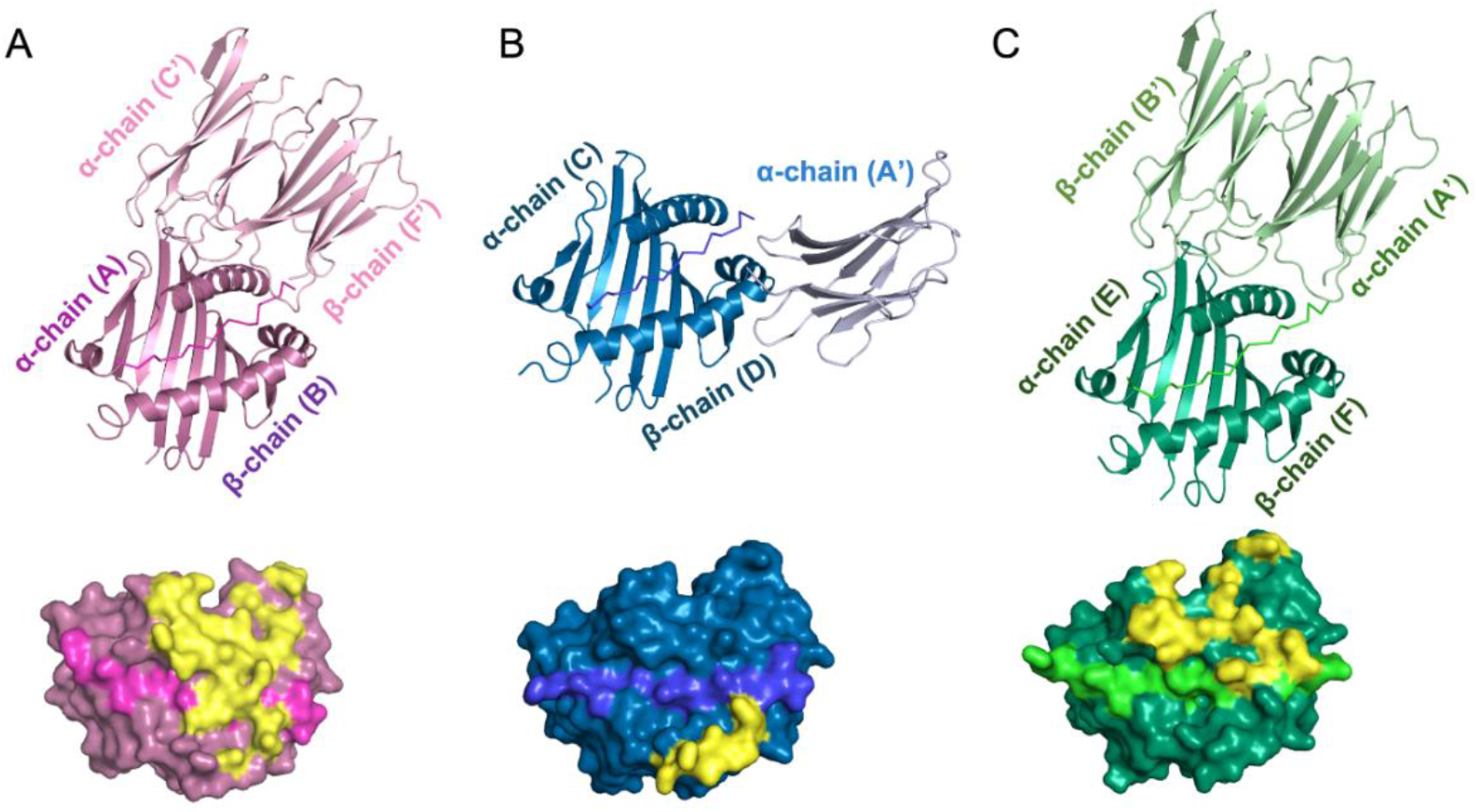
The crystal contacts from molecules outside of the asymmetric unit are different for each of the three monomers of BL2*21:01 within the asymmetric unit. Above, PBG and bound peptide of a monomer within the asymmetric unit interacting with Ig-like domains from other monomers outside of the asymmetric unit. Below, surface of PBG and bound peptide of a monomer within the asymmetric unit with footprint of interacting Ig-like domains indicated in yellow. A. A/B monomer (PyMol cartoon mode, cb_magenta) and peptide (ribbon, light magenta) interacting with chains C’ and F’ (light pink). B. C/D (cb_blue) and peptide (tv_blue) interacting with chain A’ (light blue). C. E/F (cb_green) and peptide (tv_green) interacting with chains A’ and B’ (light green).

Comparing the interactions of the adjacent Ig-like domains for the A/B and E/F monomers in more detail (Fig. 7B, Table S4), some crystal contacts between the α chains are the same, including Aspα20, Alaα22, Leuα40, Aspα41 and Lysα71 in A/B and E/F with Gluα134, Glnα160, Argα161 and Aspα163 in C’ and A’, respectively. In addition, one salt bridge is the same for both monomers (Lysα71 with Aspα163), while another salt bridge is not (Lysα71 with Aspα163 for A/B, but replaced by Gluα20 to Argα127 in in E/F). One residue, Glnα72, has multiple crystal contacts for A/B but only one for E/F; in any case it is located away from the PBG and does not contact peptide. Similarly, Asnβ64, Gluβ66 and Gluβ69 from C/D (but not from A/B) have crystal contacts but do not contact peptide. Tyrβ60 from A/B forms H-bonds with Cysβ169, Glnβ134 and Glyβ168 in F’, while Tyrβ60 from E/F does not have crystal contacts. The crystal contacts had no effect on amino acid backbones of the MHC molecules, based on superimposition of the Ca trace.

However, Argβ166 from the adjacent Ig-like domains does interact differently with the peptide in the A/B and E/F monomers. The Argβ166 from the B’ chain forms an H-bond with the mainchain carbonyl after peptide residue Leu9 of the E/F monomer with a nonamer core sequence (Fig. 7Bii), whereas the Argβ166 from the F’ chain forms an H-bond with the mainchain carbonyl after peptide residue Ala6 in the A/B monomer with the decamer core sequence (Fig. 7Bi). Having said that, the location and orientation of the carbonyl after peptide Ala6 is not significantly different (Fig. 7Bi versus ii) although the orientation of sidechain is slightly different (Fig. S7), so it seems unlikely that the interaction with Argβ166 is responsible for the difference in peptide mainchain conformation between A/B and C/D. Taking all of this evidence together, there is no strong evidence for the crystal contacts either directly or indirectly causing changes in the conformation of the bound peptide.

## Discussion

In this paper, the third structure of a chicken MHC class II molecule is reported, with the aim of evaluating whether this BL2*021:01 molecule from the MDV-resistant haplotype B21 binds peptides with the same unprecedented mode as BL2*002:01 from another MDV-resistant haplotype B2 (14), or with the typical mode as BL2*019:01 from the MDV-susceptible haplotype B19 (22). To our surprise, we discovered that BL2*021:01 binds an important MDV peptide in both modes. We speculate that binding the same peptide in two modes will increase the molecular surfaces of peptides presented by this chicken class II molecule, expand the number of T cell clones that respond to a single peptide, and thus potentially provide a greater immune response to a pathogen. Such a property may by one reason why the B21 haplotype is found so frequently among chicken flocks and populations.

Outside of typical mammals like humans and mice, the chicken is the only species for which the immune molecules have been extensively studied at the biochemical and cell biological level, so that the genetics of resistance to the many common infectious pathogens could be examined in molecular detail (38, 39). Such studies build on decades of biological research driven by the economic importance of the global poultry industry, for which huge populations of domestic chickens are monitored, analogous to outbred human populations being monitored by the public health system, followed by intense research activity and eventual intervention (40). Among the many discoveries from studying chickens, a key point is that the chicken MHC is small and simple compared to typical mammals, expressing single dominantly-expressed class I and class II molecules whose properties determine the immune response. It is not too much to say that chickens can live or die depending on their MHC haplotype, with this genomic simplicity allowing discovery of general principles that were not easy to discern from the large and complex MHC of typical mammals (2, 40).

While there have been reports of MHC class II molecules presenting peptides in different conformations (36, 41–45), none is like the example reported here. For example, class II-invariant chain peptide (CLIP) bound to class II molecules have been described to be in different sequence registers or even reversed in orientation in the PBG, and machine learning has identified significant percentages of self-peptides bound to class II molecules in reverse orientation (36, 43). However, it was not easy to find examples in the scientific literature of the exact same peptide in the same register having different conformations in any MHC molecule. Almost all MHC structures reported are from typical mammals, which may represent only a subset of the possibilities (21). Indeed, having a single dominantly-expressed class I or class II molecule as in chickens may mean that the selection pressure for plasticity is much stronger than on any one member of a multigene family as in typical mammals (3). By displaying the same peptide in different conformations, a single class II molecule may stimulate a protective T cell response with a similar efficacy as a multigene family, with important ramifications for disease resistance and vaccine responses.

## Acknowledgements

We thank Maria Danysz for technical assistance, Dr. Paul Brear of the Crystallography facility in the Department of Biochemistry at the University of Cambridge for much help, and the staff of the beamline I04 at the Diamond Light Source for help with data collection. We thank the Wellcome Trust (Investigator Award 110106/A/15/Z to JK, Senior Research Fellowship 200898 to AGC), the Biotechnology and Biological Sciences Research Council (BBSRC project grants BB/V000756/1 to JK, BB/T020954/2 to SB), the German Science Foundation (Deutsche Forschungsgemeinschaft FOR5130, KA 5564/1-1 to JK) and the University of Edinburgh Immunology Senior Honours program to JK. We thank Matthew Spencer for critical reading. For open access, the author has applied a Creative Commons Attribution (CC BY) licence to any Author Accepted Manuscript version arising from this submission.

