## Supplementary Information for "A major histocompatibility complex (MHC) class II molecule that binds the same viral pathogen peptide with both nonamer and decamer core sequences for presentation to T cells"

**Supplementary table S1.** Data collection and refinement statistics (https://www.rcsb.org/structure/7PDY). Values in parentheses are for highest resolution shell.

**Supplementary table S2.** Hydrogen bonds formed between the peptide and the class II (BL2*21:01) molecule within each of the monomers. Amino acids of the peptides are numbered from the first anchor residue; negative numbers indicate the N-terminal flanking region. Amino acids of the class II molecule numbered from the start of the mature protein. Bond lengths were measured in Å as a distance between the atoms forming hydrogen bonds. No salt bridges were detected between the peptide residues and the class II molecules.

**Supplementary table S3.** The hydrogen bonds and salt bridges formed between the peptide of each monomer with the other class II (BL2*21:01) monomers within the asymmetric unit. Amino acids of the class II molecule numbered from the start of the mature protein. Bond lengths were measured in Å as a distance between the atoms forming hydrogen bonds or salt bridges (with salt bridges in red).

**Supplementary table S4.** The hydrogen bonds and salt bridges formed between the monomers within the asymmetric unit of BL2*021:01 with the molecules outside the asymmetric unit (that is, the crystal contacts). Amino acids of the class II molecule numbered from the start of the mature protein. Bond lengths were measured in Å as a distance between the atoms forming hydrogen bonds or salt bridges (with salt bridges in red).

**Supplementary table S1**

| | 7PDY |
|---|---|
| **Data Collection** | |
| Wavelength (Å) | 0.920 |
| Resolution range (Å)<br>(highest resolution shell) | 103.3 - 2.543<br>(2.634 - 2.543) |
| Space group | $P\ 3_2\ 2\ 1$ |
| Unit cell<br>a,b,c, (Å)<br>α,β,γ (°) | 238.54, 238.54,<br>76.52<br>90, 90, 120 |
| Total reflections | 1686668 |
| Unique reflections | 81217 (8140) |
| Multiplicity | 20.7 |
| Completeness (%) | 99.02 (99.84) |
| Mean I/sigma (I) | 16.6 (0.8) |
| Wilson B-factor (Å$^2$) | 75.43 |
| R-merge | 0.14 (4.4) |
| R-meas | 0.14 (4.5) |
| R-pim | 0.03 (0.99) |
| CC1/2 | 0.99 (0.44) |
| **Refinement** | |
| Reflections used in refinement | 77013 |
| Reflections used for R-free | 4205 |
| $R_{work}$ | 0.22 |
| $R_{free}$ | 0.25 |
| Number of atoms:<br>non-hydrogen | 9525 |
| macromolecules | 9224 |
| Ligands | 154 |
| Solvent | 147 |
| Protein residues | 1165 |
| RMS (bonds, Å) | 0.007 |
| RMS (angles, °) | 1.422 |
| Ramachandran favored (%) | 98 |
| Ramachandran allowed (%) | 2 |
| Ramachandran outliers (%) | 0 |
| Rotamer outliers (%) | 4.62 |
| Clash score | 2.65 |
| Average B-factor (Å$^2$) | 83.75 |
| Macromolecules | 83.11 |
| Ligands | 135.68 |
| Solvent | 69.31 |
| Number of TLS groups | 6 |

### Supplementary table S2

| Peptide bound to chains: A/B | | | Peptide bound to chains: C/D | | | Peptide bound to chains: E/F | | |
|---|---|---|---|---|---|---|---|---|
| Peptide | Distance (Å) | Chain A (α-chain) | Peptide | Distance (Å) | Chain C (α-chain) | Peptide | Distance (Å) | Chain E (α-chain) |
| | | | | | | P -3: [O] | 3.72 | S57: [OG] |
| A-2: [O] | 2.90 | S57: [N] | A-2: [O] | 2.83 | S57: [N] | A-2: [O] | 2.86 | S57: [N] |
| A-2: [O] | 3.48 | S57: [OG] | A-2: [O] | 2.81 | S57: [OG] | A-2: [O] | 3.44 | S57: [OG] |
| A-2: [N] | 3.86 | F55: [O] | | | | | | |
| V1: [N] | 2.82 | S57: [O] | V1: [N] | 2.86 | S57: [O] | V1: [N] | 2.89 | S57: [O] |
| V4: [N] | 2.84 | Q9: [OE1] | V4: [N] | 2.72 | Q9: [OE1] | V4: [N] | 2.81 | Q9: [OE1] |
| V4: [O] | 2.93 | Q9: [NE2] | V4: [O] | 3.44 | Q9: [NE2] | V4: [O] | 3.04 | Q9: [NE2] |
| V4: [O] | 3.02 | N66: [ND2] | V4: [O] | 2.71 | N66: [ND2] | V4: [O] | 3.17 | N66: [ND2] |
| A6: [N] | 3.47 | N66: [OD1] | | | | | | |
| | | | L7: [O] | 2.83 | N73: [ND2] | L7: [O] | 3.03 | N73: [ND2] |
| | | | L9: [N] | 3.12 | N73: [OD1] | L9: [N] | 2.92 | N73: [OD1] |
| A10: [N] | 2.63 | N73: [OD1] | A10: [O] | 3.31 | N80: [ND2] | A10: [O] | 3.07 | N80: [ND2] |
| E11: [O] | 2.91 | N80: [ND2] | | | | | | |
| | | | R12: [N] | 3.17 | N80: [OD1] | R12: [N] | 3.23 | N80: [OD1] |
| Q13: [N] | 2.98 | N80: [OD1] | | | | | | |
| Peptide | Distance (Å) | Chain B (β-chain) | Peptide | Distance (Å) | Chain D (β-chain) | Peptide | Distance (Å) | Chain F (β-chain) |
| V-1: [O] | 2.69 | H81: [NE2] | | | | V-1: [O] | 3.00 | H81: [NE2] |
| H2: [N] | 2.72 | N82: [OD1] | H2: [N] | 3.15 | N82: [OD1] | H2: [N] | 3.03 | N82: [OD1] |
| H2: [O] | 2.75 | N82: [ND2] | H2: [O] | 3.23 | N82: [ND2] | H2: [O] | 2.80 | N82: [ND2] |
| H2: [ND1] | 3.21 | R77: [O] | H2: [ND1] | 3.02 | R77: [O] | | | |
| S3: [O] | 2.61 | R77: [NH1] | S3: [O] | 3.20 | R77: [NH1] | S3: [O] | 2.69 | R77: [NH1] |
| R5: [N] | 3.73 | E74: [OE2] | R5: [N] | 3.63 | E74: [OE2] | R5: [N] | 3.78 | E74: [OE2] |
| | | | R5: [O] | 3.63 | K11: [NZ] | | | |
| | | | M8: [O] | 3.07 | W61: [NE1] | M8: [O] | 2.87 | W61: [NE1] |
| L9: [O] | 3.05 | W61: [NE1] | | | | | | |
| | | | A10: [N] | 3.19 | Q57: [OE1] | A10: [N] | 2.85 | Q57: [OE1] |
| | | | A10: [O] | 3.31 | Q57: [NE2] | A10: [O] | 3.21 | Q57: [NE2] |
| E11: [N] | 2.91 | Q57: [OE1] | | | | | | |
| E11: [O] | 3.22 | Q57: [NE2] | | | | | | |

### Supplementary table S3

| Peptide bound to chains: A/B | | | Peptide bound to chains: C/D | | | Peptide bound to chains: E/F | | |
|---|---|---|---|---|---|---|---|---|
| Chain C (α-chain) | Distance (Å) | Chain F (β-chain) | Chain C (α-chain) | Distance (Å) | Chain E (α-chain) | Chain D (β-chain) | Distance (Å) | Chain E (α-chain) |
| Q128: [O] | 2.88 | R141: [NH1] | E92: [OE1] | 2.79 | K115: [NZ] | R34: [NH2] | 2.92 | D146: [OD1] |
| | | | | | | R34: [NH1] | 3.11 | D146: [OD2] |
| G129: [N] | 3.66 | E162: [OE1] | K115: [NZ] | 2.88 | E92: [OE1] | R34: [NH1] | 3.65 | D146: [OD1] |
| | | | | | | R34: [NH2] | 3.07 | D146: [OD2] |
| D146: [OD2] | 2.85 | R34: [NH1] | | | | | | |
| D146: [OD1] | 2.78 | R34: [NH1] | | | | G107: [O] | 2.73 | V186: [N] |
| D146: [OD2] | 3.53 | R34: [NH2] | | | | G107: [N] | 3.33 | E183: [OE2] |
| R161: [NE] | 2.76 | S108: [O] | | | | S108: [OG] | 2.67 | G162: [N] |
| | | | | | | S108: [O] | 2.83 | R161: [NE] |
| G162: [N] | 2.85 | S108: [OG] | | | | S108: [N] | 3.81 | P184: [O] |
| Q179: [OE1] | 2.83 | S144: [N] | | | | R114: [NE] | 3.01 | E183: [OE1] |
| Q179: [NE2] | 3.61 | V142: [O] | | | | R114: [NH2] | 3.55 | E183: [OE2] |
| | | | | | | R114: [NH1] | 3.98 | E183: [OE1] |
| R180: [NH2] | 3.13 | D146: [OD2] | | | | R114: [NH2] | 3.63 | E183: [OE1] |
| D183: [OE2] | 2.86 | G107: [N] | | | | E140: [OE1] | 3.84 | R126: [NH2] |
| E183: [OE1] | 3.30 | R114: [NH1] | | | | E140: [OE1] | 3.84 | R126: [NH2] |
| E183: [OE1] | 3.65 | R114: [NE] | | | | E140: [OE1] | 3.73 | R126: [NH1] |
| E183: [OE2] | 3.75 | R114: [NH1] | | | | | | |
| | | | | | | R141: [NH2] | 3.17 | N128: [O] |
| V186: [N] | 2.62 | G107: [O] | | | | | | |

### Supplementary table S4

| Molecule | Residue / Main Chain-Number: [atom] | Distance (Å) | Residue / Symm. Chain-Number: [atom] |
|---|---|---|---|
| Peptide bound to chains A/B | peptide A6: [O] | 3.19 | RβF’166: [NE] |
| | peptide E11: [OE1] | 2.76 | NβF’134: [ND2] |
| Chains A/B | QαA72: [NE2] | 3.73 | TβF’163: [OG1] |
| | QαA72: [NE2] | 3.75 | VβF’164: [O] |
| | QαA72: [OE1] | 3.74 | TβF’163: [OG1] |
| | QαA72: [OE1] | 2.93 | VβF’164: [N] |
| | DαA20: [N] | 3.15 | EαC’134: [OE1] |
| | KαA71: [NZ] | 2.68 | RαC’161: [O] |
| | KαA71: [NZ] | 2.70 | DαC’163: [OD1] |
| | AαA22: [O] | 3.14 | QαC’160: [ NE2] |
| | DαA41: [O] | 2.34 | RαC’161: [NH1] |
| | LαA40: [O] | 2.72 | RαC’161: [NH1] |
| | KαA71: [NZ] | 3.90 | DαC’163: [OD2] |
| | KαA71: [NZ] | 2.70 | DαC’163: [OD1] |
| | YβB60: [OH] | 2.51 | DβF’169: [OD1] |
| | YβB60: [OH] | 3.56 | NβF’134: [OD1] |
| | YβB60: [OH] | 3.53 | GβF’168: [O] |
| Chains C/D | NβD64: [ND2] | 3.23 | PβD’110: [O] |
| | EβD66: [OE1] | 3.51 | TβD’112: [N] |
| | EβD66: [OE1] | 3.33 | TβD’112: [OG1] |
| | EβD66: [OE2] | 3.15 | TβD’112: [OG1] |
| | EβD69: [OE1] | 3.74 | RβD’167: [N] |
| | EβD69: [OE2] | 3.44 | RβD’167: [N] |
| Peptide bound to chains E/F | peptide L9: [O] | 3.63 | RβB’166: [NH2] |
| Chains E/F | DαE20: [N] | 3.17 | EαA’134: [OE1] |
| | KαE71: [NZ] | 3.18 | DαA’163: [OD1] |
| | DαE20: [OD2] | 2.75 | RαA’127: [NH1] |
| | AαE22: [O] | 2.86 | QαA’160: [NE2] |
| | AαE40: [O] | 2.59 | RαA’161: [NH1] |
| | DαE41: [O] | 2.82 | RαA’161: [NH1] |
| | KαE71: [NZ] | 3.18 | DαA’163: [OD1] |
| | DαE20: [OD2] | 2.75 | RαA’127: [NH1] |

**>BLA_construct**

MLLVNQSHQGFNKEHTSKMVSAIVLYVLLAAAAHSAFAADRR**HVLLQAEFYQRSEGPDKAWA**
**QFGFHFDADELFHVELDAAQTVWRLPEFGRFASFEAQGALQNMAVGKQNLEVMISNSNR**
**SQQDFVTPELALFPAEAVSLEEPNVLICYADKFWPPVATMEWRRNGAVVSEGVYDSVYYGR**
**PDLLFRKFSYLPFVPQRGDVYSCAVRHWGAEGPVQRMW**GGGSRSGGGSTDTLQAETDQL
EDEKSALQTEIANLLKEKEKLEFILAAYGGGSGGGSGTGLNDIFEAQKIEWHE-

**>BL2*021-pp38_construct**

MLLVNQSHQGFNKEHTSKMVSAIVLYVLLAAAAHSAFAADRP*AVVHSVRALMLAERQ*GGGGS
GGGGSGGGG**SFFYGKIGECHYLNGTERVRFLDRQIYNRQQFAHFDSDVGKFVADTPLGEP**
**QAEYWNSNAELLENLMNEVDRVCRHNYGILESFTVQRSVEPKVRVSALQSGSLPETDRLA**
**CYVTGFYPPEIEVKWFLNGREETERVVSTDVMQNGDWTYQVLVVLETVPRRGDSYVCRVE**
**HASLRQPISQAWEPPADAGRSK**GENLYFQGGSIARLEEKVKTLKAQNSELASTANMLREQVA
QLKQKVMNGGGSGGGSSRGPFEGKPIPNPLLGLDSTRTG**HHHHHH**-

Signal peptide, linkers, Avi-tag, c-fos, jun, **MHC II**, *peptide*, TEV protease cleavage site,
V5-tag, **His-tag**.

Supplementary Figure S1. Amino acid sequences of the BLA and pp38 peptide-BLB2*21:01 constructs used in this study. Single letter amino acid code; tags and linkers are as coloured.

A

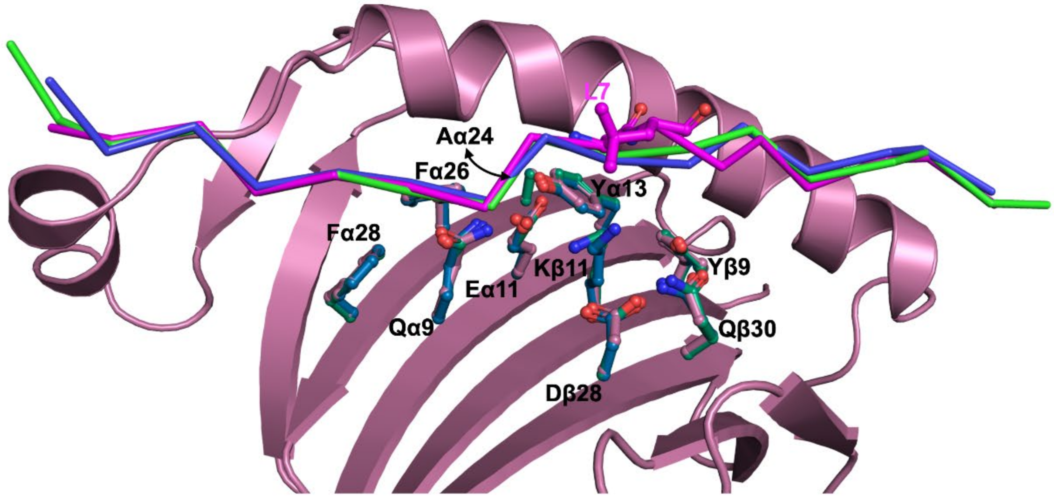

B

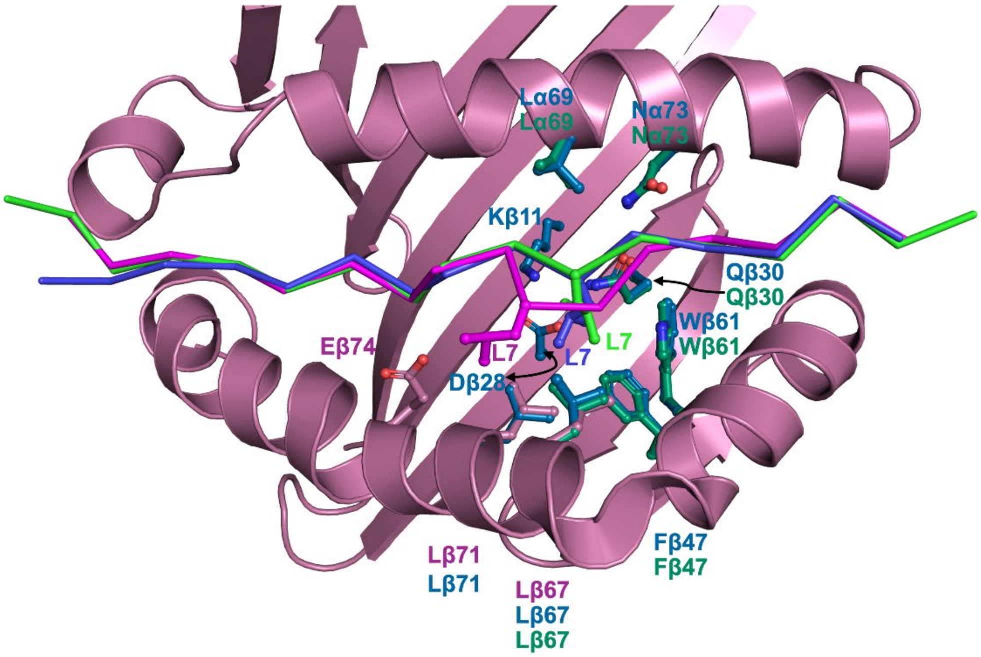

Supplementary Figure S2. Amino acids of PBG that are in close proximity to the peptide Leu7 (L7).
A. Amino acid side chains shown in sticks for all residues on the β-strands of the PBG under L7 that are pointing upwards (side chains that are pointing downwards were excluded). B. Amino acids that are within 4 Å of L7 of the peptide bound to chains A/B (cb_violet), chains C/D (cb_blue), and chains E/F (cb-green) are shown in sticks and labelled accordingly. Those amino acids with distances to the peptide position L7 of more than 4 Å are not depicted.

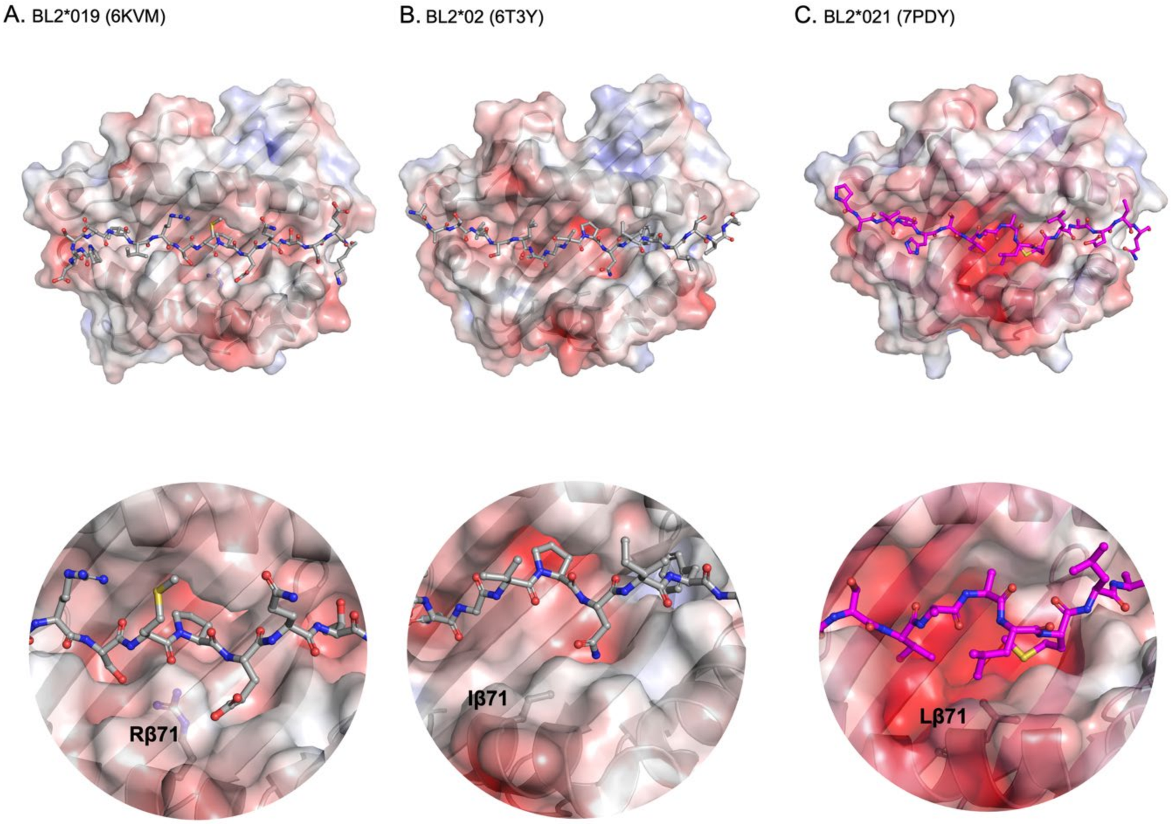

Supplementary Figure S3. Comparison of the surface of the PBG of three chicken MHC molecules.
A. Argβ71 (Rβ71) of BL2*019 (6KVM) forms a bulge in the PBG narrowing it down, in comparison to B. Ileβ71 (Iβ71) of BL2*02 (6T3Y) and C. Lβ71 (Lβ71) of BL2*021 (7PDY). The peptides are presented in sticks.

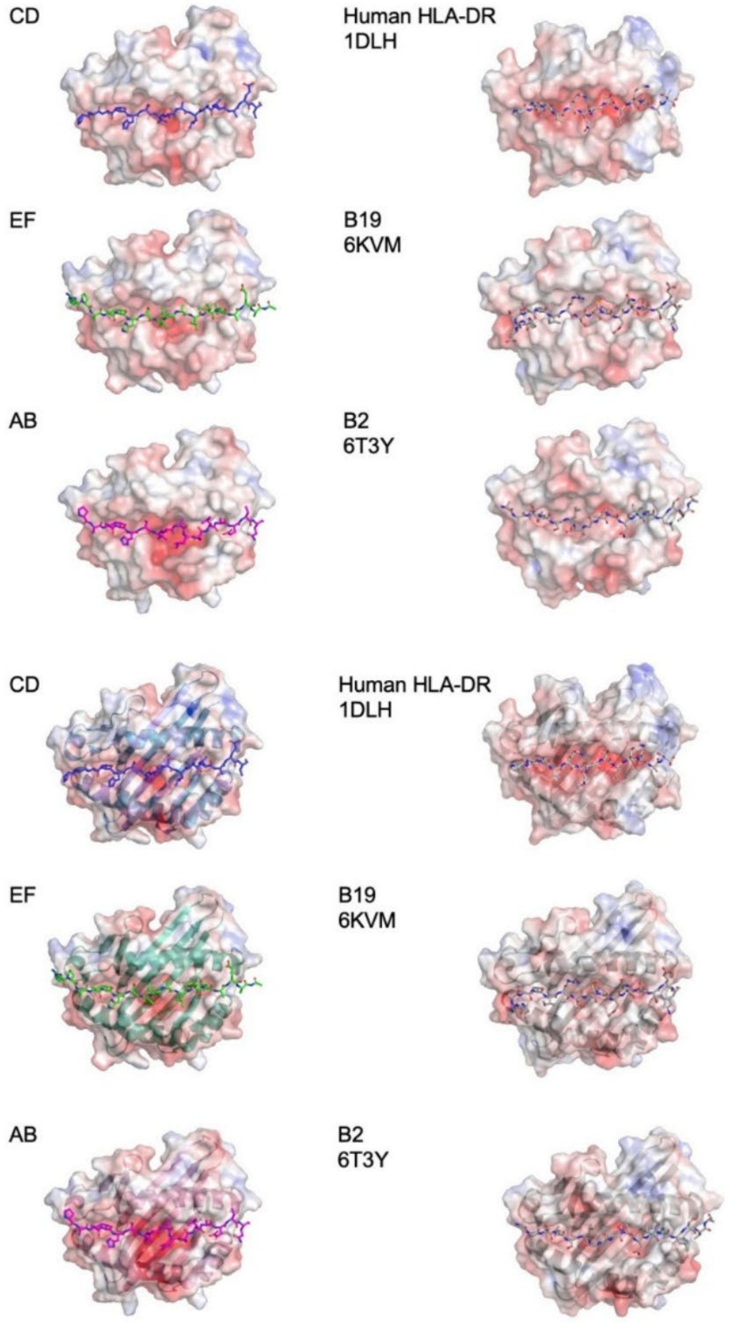

Supplementary Figure S4. Top view of the peptide-binding groove (PBG) from MHC class II molecules analysed in this study. Top six panels: solvent-accessible surface of class II molecules calculated by APBS electrostatics (positive charge, blue; negative charge, red) with peptides in sticks. Bottom six panels, the same samples as above, but with the surface view set to 40% transparency view the Cα backbone of MHC class II molecules. AB, CD and EG, three monomers within the asymmetric unit (7PDY).

A

B

C

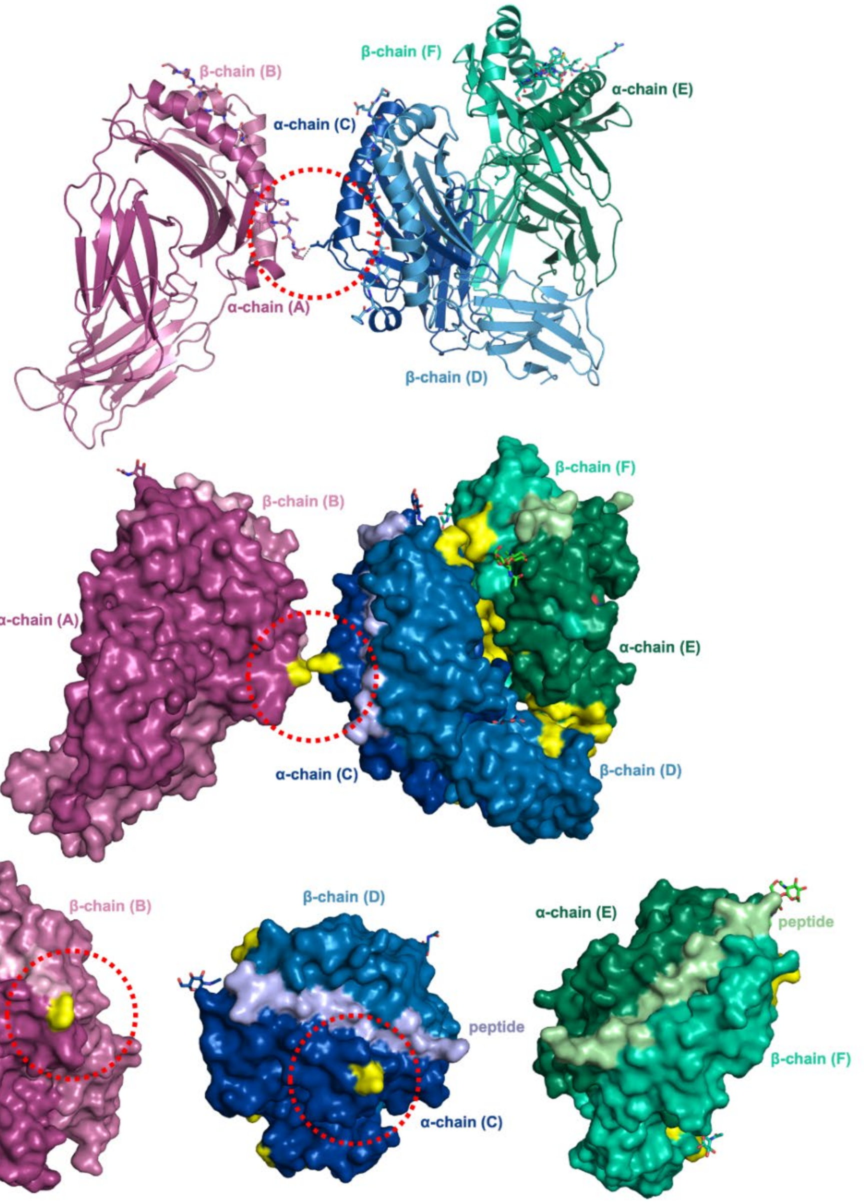

Supplementary Figure S5. Footprints of the three BL2*21:01 molecules in the asymmetric unit with respect to each other. A. Proline ($P_{-3}$) of the peptide bound to chains A/B (cartoon, cb-violet) is the only residue of all three peptides bound to BL2*21:01 that is within 4 Å of chains C/D (cartoon, cb_blue) highlighted by the red dotted circle. Note that $P_{-3}$ does not from any hydrogen bond with amino acid residues from chains C/D. B. Surface representation of the three BL2*21:01 molecules in the asymmetric unit where the footprints of each BL2*21:01 molecule were determined by their distance (within 4 Å) to the adjacent molecule and indicated in yellow.  C. Looking from top of the PBG of each of the three BL2*21:01 molecules in the asymmetric unit, to show that the only footprint that includes a peptide is that bound to chains A/B (light pink). The two peptides bound to chains C/D (light blue) and E/F (light green) do not form any contact with any adjacent chain.

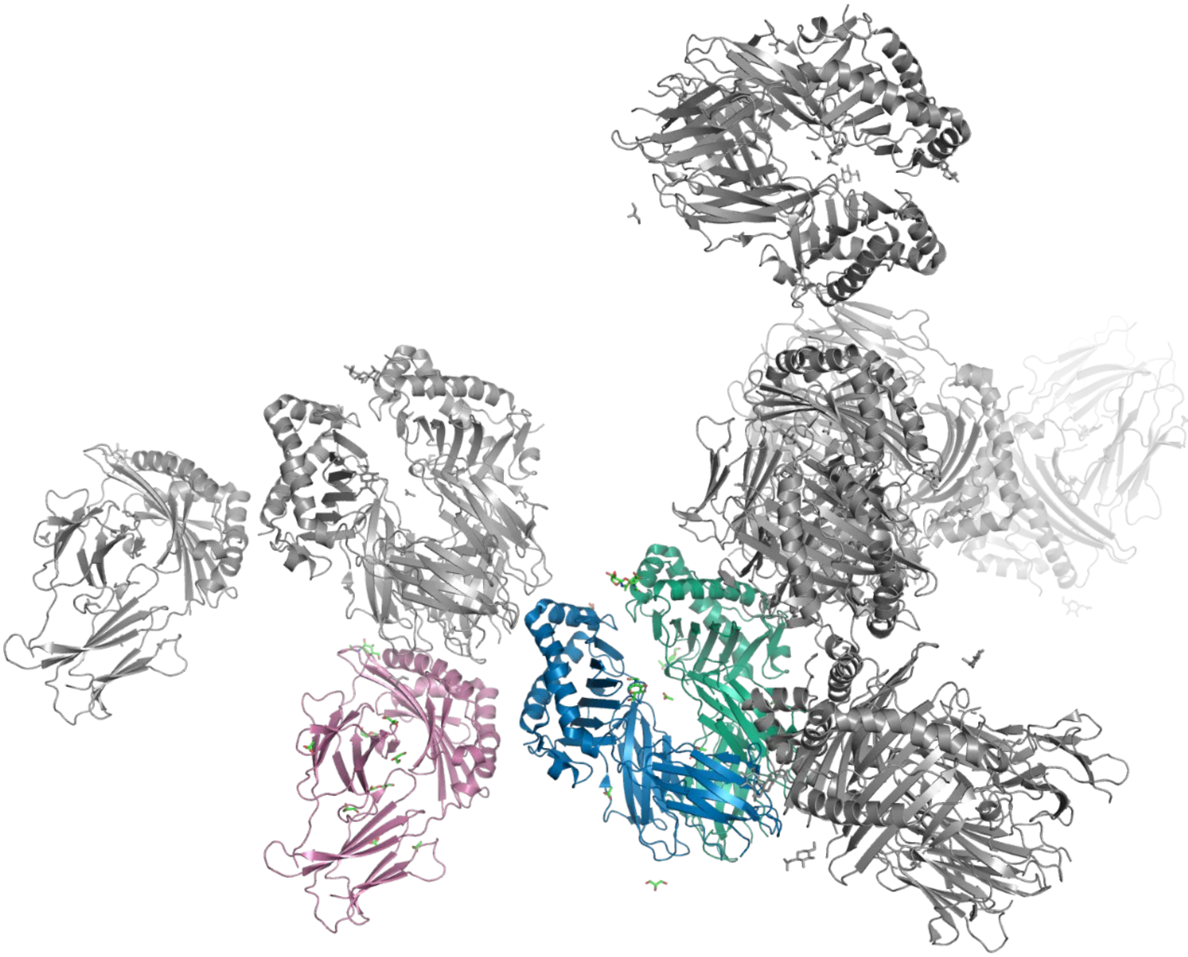

Supplementary Figure S6. The peptide-binding groove (PBG) of the three monomers of peptide-BL2*021:01 in the asymmetric unit have different types of crystal contacts. Class II molecules in PyMol cartoon mode and peptide in PyMol sticks. Monomers in asymmetric unit are A/B in magenta, C/D in blue and E/F in green; monomers outside of the asymmetric unit are in grey.

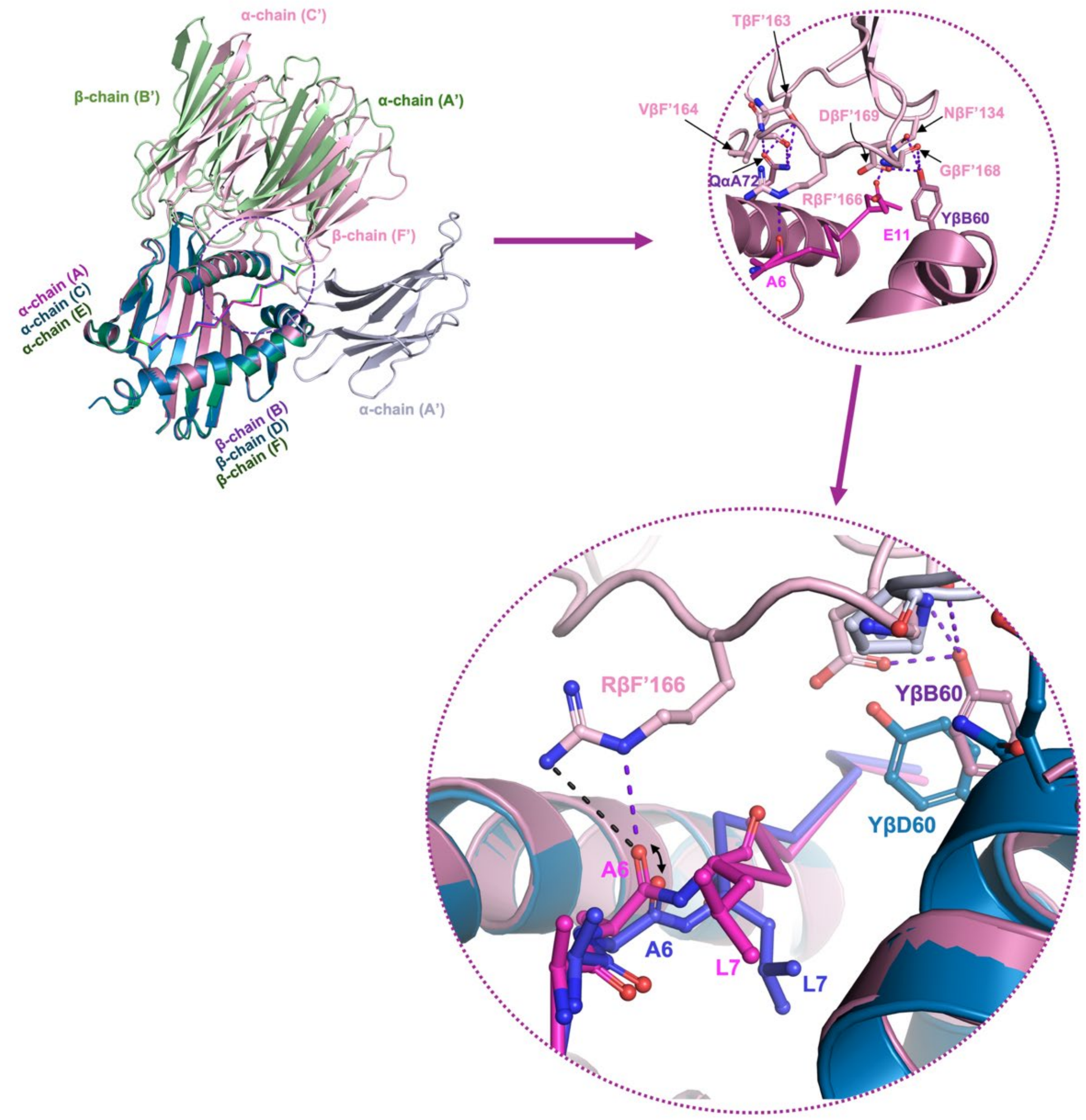

Supplementary Figure S7. Alanine at position 6 (A6) of the peptide adopts a slightly different spatial orientation when bound to chains A/B (cb_violet), compared to the conformation in the peptide bound to chains C/D (cb_blue). The diverging of the carbonyl of A6 is indicated by an arrow in the bottom panel. Note that A6 of chains A/B form a hydrogen bond with Arg from the β chain of the F' chain (RβF'166, light pink) as part of the crystal contact with the chain F' of the adjacent asymmetric unit.
